# Speedy benthi: Elongated photoperiods reduce generation times of the model plant species *Nicotiana benthamiana*

**DOI:** 10.1101/2024.01.18.576090

**Authors:** Matthew Castle, Daniel Lüdke

## Abstract

*Nicotiana benthamiana* is increasingly gaining prominence as a model plant species with recently published high-quality genome assemblies, which will further enable forward and reverse genetic approaches (Bally et al., 2018; Derevnina et al., 2019; Kourelis et al., 2019; Ranawaka et al., 2023; Vollheyde et al., 2023). However, the generation time of *N. benthamiana* poses a bottleneck in the creation of mutant and transgenic plant lines. Speed breeding (SB), by extended photoperiods and adjustments to growth parameters, is an efficient way to reduce generation times for many crop and model plant species (Ghosh et al., 2018; Watson et al., 2018; Hickey et al., 2019; Varshney et al., 2021). We hypothesized that an extended photoperiod could reduce the seed to seed generation time of *N. benthamiana*. We tested this hypothesis by comparing generation times under SB conditions to traditionally used photoperiods in growth chambers and green house settings. We found that a 22h photoperiod reduced the generation time of *N. benthamiana* by approximately 2 weeks (16-22%). Fertilization in combination with a far-red light spectrum did not yield a further reduction in generation time when combined with SB conditions. Our findings further contribute to the establishment of *N. benthamiana* as an important model organism for plant research.

## Results

We performed seed to seed generation growth trials with extended photoperiods in controlled growth chambers (Supplemental Data 1). Based on established methods for breeding crop and model plant species, a 22h photoperiod with 2h of darkness was used as SB condition (Ghosh et al., 2018; Watson et al., 2018). The control condition for normal breeding (NB) consisted of a 16h photoperiod and 8h of darkness (Ghosh et al., 2018; Watson et al., 2018). The seed to seed generation time was determined from seed planting on soil (Levington F2 Starter) to opening of the first seed pod. For both, SB and NB conditions in controlled growth chambers, a temperature of 22°C and 80% humidity was used. As an additional control, we planted seeds on soil and grew them in a greenhouse facility with seasonal light, temperature, and humidity during the 2023 Norwich (UK) summer months (mid-June to August).

Growing *N. benthamiana* under SB conditions resulted in a mean seed to seed generation time of 67.65 ± 2.13 days post planting (dpp), while the NB and GH condition resulted in generation times of 78.76 ± 2.76 and 82.75 ± 1.25 dpp, respectively (Figure 1B). This constitutes a mean reduction of 11 days (16.42%) for the SB compared to the NB condition, and 15 days (22.32%) compared to the GH condition (Figure 1B, Supplemental Data 2). Notably, the extended SB photoperiod resulted in slightly enhanced plant height and yellowing symptoms, possibly due to increased light stress or nutrient deficiencies (Figure 1A, Supplemental Data 2).

**Figure 1.**
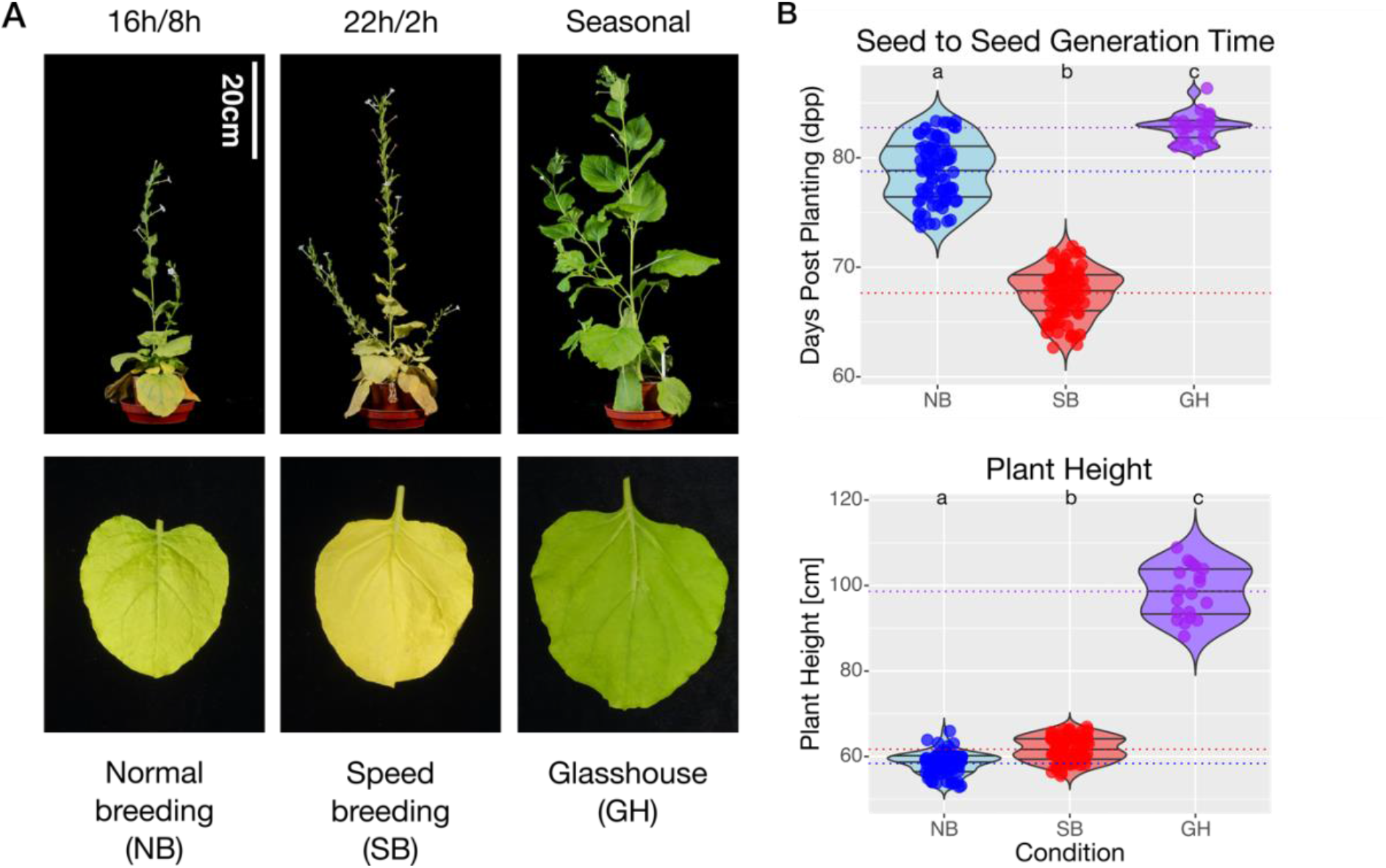
Extended photoperiods reduce the generation time of *Nicotiana benthamiana*. **A**, Representative pictures of 60 day old plants (top) and lower leaves (bottom) of *N. benthamiana* grown under normal breeding (NB), speed breeding (SB), or glasshouse (GH) conditions. **B**, Violin plots showing seed to seed generation times (top) of *N. benthamiana* and the plant height at the timepoint of the first seed pod opening (bottom). Plants were grown under normal breeding (NB), speed breeding (SB), or glasshouse (GH) conditions. The dotted lines indicate the mean values, 0.25, 0.5, and 0.75 quantiles are indicated by black lines within the respective violin plot. Normal distribution of the data was assessed using the Shapiro-Wilk-test, the Kruskal-Wallis test followed by Dunn’s posthoc test with Bonferroni correction was used to determine significant differences between the conditions (P < 0.05). Significant differences are indicated by different letters above the violin plot. All code and data are available under https://github.com/danlue/Speedy-benthi (Lüdke, 2024).

Other speed breeding approaches used fertilization to counteract possible stress responses and/or nutrient deficiencies (Ghosh et al., 2018). Moreover, the use of a far-red light spectrum was shown to additionally reduce the generation time for some crop plants (Ghosh et al., 2018; Baguma et al., 2023). We speculated that fertilization (Supplemental Data 3) could alleviate the yellowing phenotype observed under SB conditions (Figure 1,) and in combination with the use of a far-red light spectrum (Supplemental Data 4) could further reduce the generation time of *N. benthamiana*. We therefore compared the seed to seed generation time of plants grown with fertilization (F) to plants grown without fertilization (C) under SB conditions, combined with a far-red light spectrum (R) or a normal light spectrum (N) in controlled growth chambers.

The use of fertilizer under SB conditions with a normal light spectrum (FN) slightly reduced the yellowing phenotype when compared to plants grown without fertilizer under SB conditions and normal light spectrum (CN)(Figure 2A). However, the FN condition resulted in a mean seed to seed generation time of 80.55 ± 2.61 dpp, as compared to 72.35 ± 2.21 dpp for the CN condition (Figure 2B, Supplemental Data 5). Combining fertilization with a far-red light spectrum (FR) resulted in a generation time of 70.45 ± 2.30 dpp, a marked decreased when compared to the FN condition, but insignificant when compared to the CN or the CR (68.85 ± 2.90 dpp) conditions (Figure 2B, Supplemental Data 5). We moreover observed that fertilization, and the addition of a far-red light spectrum, both in combination with and without fertilization, increased the plant height (Figure 2B). The germination rates of seeds collected from plants grown under the different conditions were not markedly different (Supplementary Figure 1). We conclude that speed breeding with an extended 22h photoperiod is sufficient to reduce the seed to seed generation time of *N. benthamiana*. Additional fertilization and/or the addition of a far-red light spectrum used in our trials do not aid in further reducing the generation time in a meaningful manner, but impact plant height.

**Figure 2.**
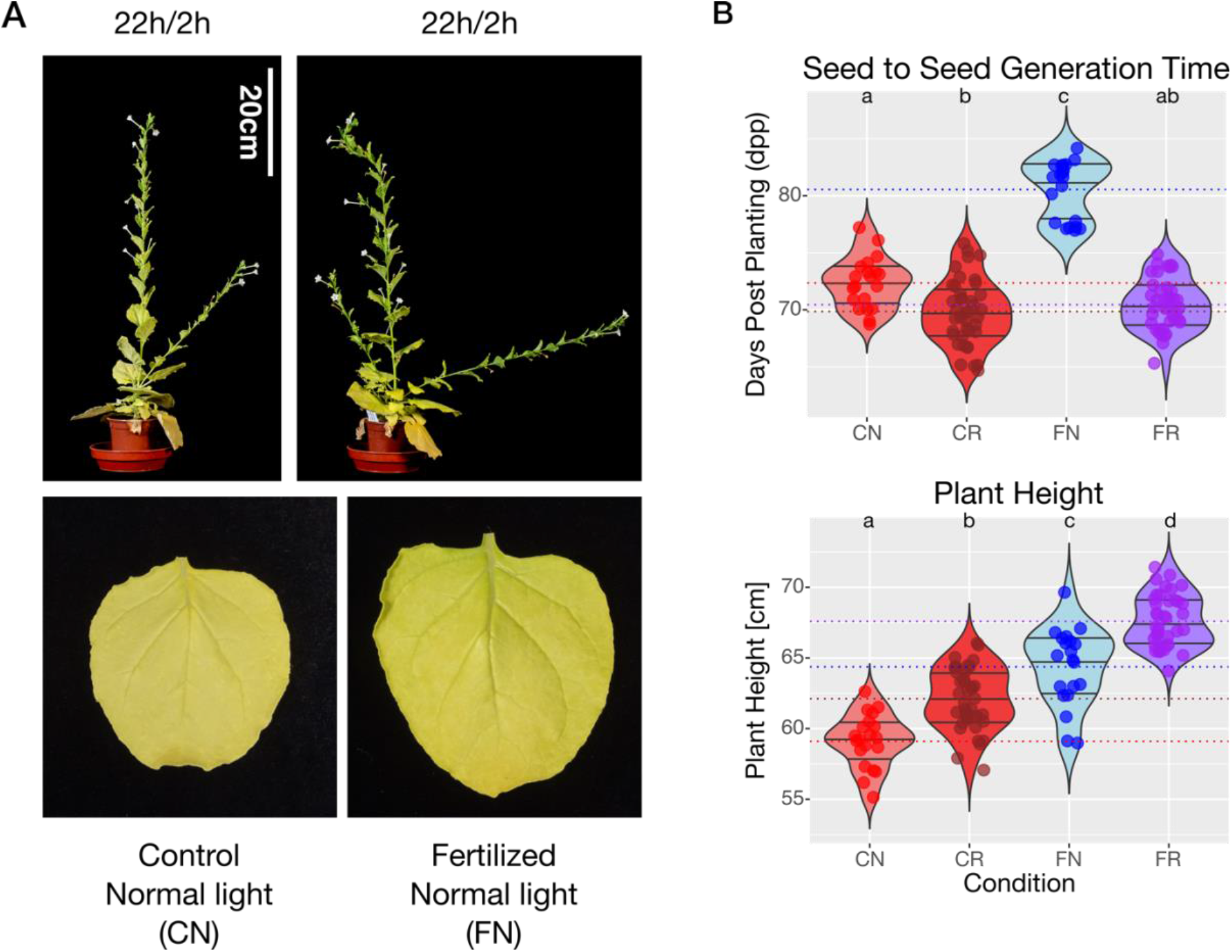
Fertilization and far-red light spectrum addition do not further reduce the generation time of *Nicotiana benthamiana* grown under extended photoperiods. **A**, Representative pictures of 60 day old plants (top) and lower leaves (bottom) of *N. benthamiana* grown under speed breeding conditions, with or without fertilization. **B**, Violin plots showing seed to seed generation times (top) of *N. benthamiana* and the plant height at the timepoint of the first seed pod opening (bottom). Plants were grown under speed breeding conditions, either without fertilizer and without far-red light addition (CN), without fertilizer and far-red light addition (CR), with fertilizer and without far-red light addition (FN), or with fertilizer and far-red light addition (FR). The dotted lines indicate the mean values, 0.25, 0.5, and 0.75 quantiles are indicated by black lines within the respective violin plot. For data on seed to seed generation times, normal distribution was assessed using the Shapiro-Wilk-test, the Kruskal-Wallis test followed by Dunn’s posthoc test with Bonferroni correction was used to determine significant differences between the conditions (P < 0.05). For data on plant height, normal distribution was assessed using the Shapiro-Wilk-test, analysis of variance was performed using Levene’s test, and one-way analysis of variance (anova) followed by Tukey’s HSD was performed to determine significant differences between the conditions (P < 0.05). Significant differences are indicated by different letters above the violin plot. All code and data are available under https://github.com/danlue/Speedy-benthi (Lüdke, 2024).

Our trial study provides evidence that a simple and cost-effective extension of the photoperiod is sufficient to reduce the seed to seed generation time of the model plant species *N. benthamiana* (Figure 1). Fertilization had a negative effect on the acceleration of the generation time without the use of far-red light treatment, but reduced the yellowing stress symptoms observed (Figure 1; Figure 2). As abiotic stress can be an additional factor to reduce the generation time of plants (Ghosh et al., 2018; Schilling et al., 2023), we speculate that additional light stress in combination with reduced nutrient availability could be a driving factor in generation time reduction in *N. benthamiana*. Considering that *N. benthamiana* originated from predominantly arid zones of Australasia, this could represent a survival strategy under naturally occurring conditions (Byrne et al., 2008; Bally et al., 2018; Schiavinato et al., 2020; Ranawaka et al., 2023). Our study further outlines that *N. benthamiana* represents a valuable tool as a model species in plant science that is amenable to generation time reduction.

**Supplementary Figure 1.**
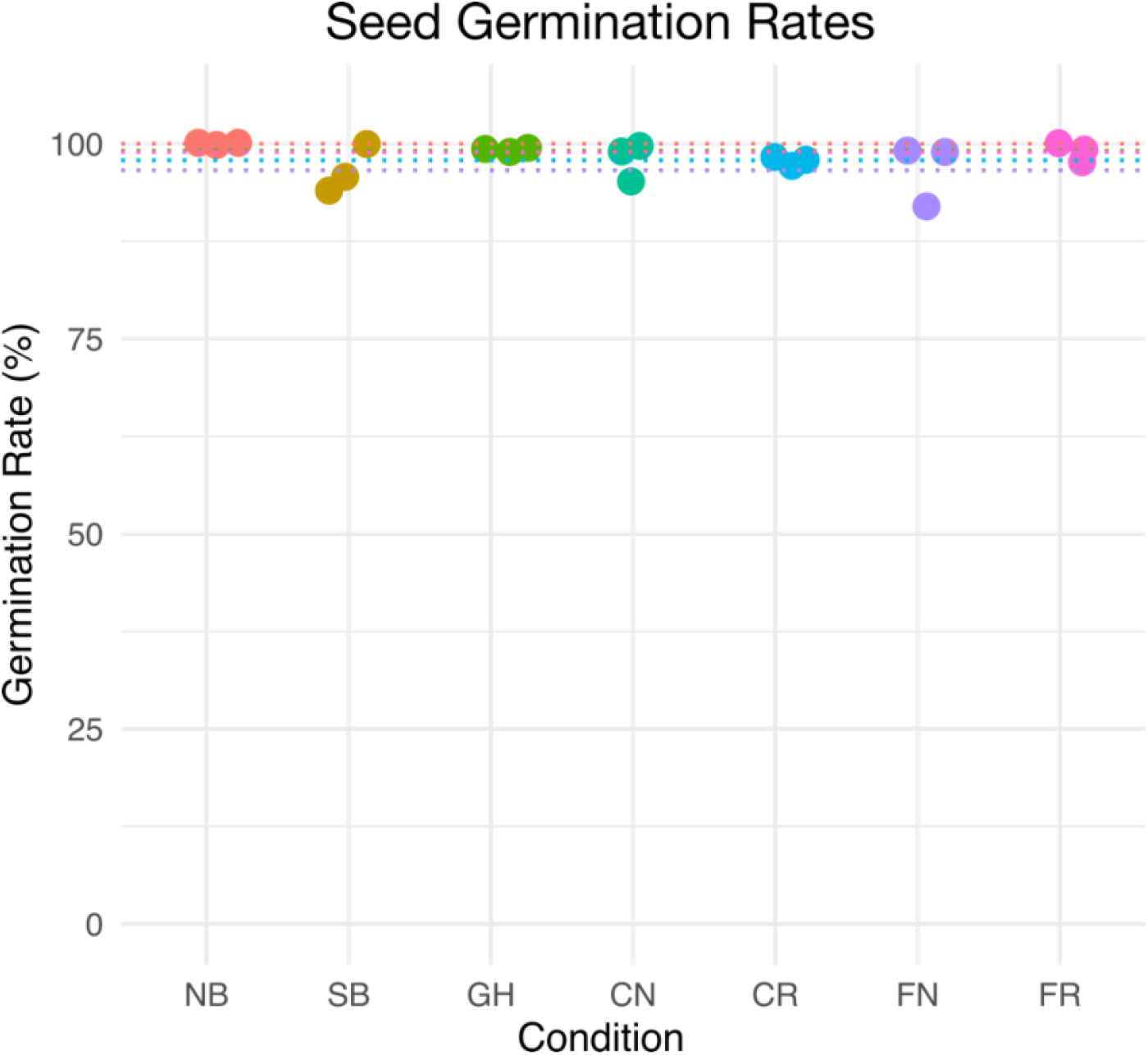
Growth conditions of *Nicotiana benthamiana* do not markedly influence the seed germination rates in the following generation. Scatter plots showing the germination rates of seeds contained in the first seed pod of *Nicotiana benthamiana* grown under the different photoperiod, fertilization and light spectrum conditions indicated in Figure 1 and 2. Seeds were sown on a paper towel (WypAll L20, Kimberly-Clark) soaked with 50ml of ¼ MS medium (1.1 g/L Murashige and Skoog medium, 0.5 g/L 2-(N-morpholino)ethanesulfonic acid, adjusted to pH 5.7) and grown for 7 days under ambient light and room temperature. All code and data are available under https://github.com/danlue/Speedy-benthi (Lüdke, 2024).

## Acknowledgment

We thank Jiorgos Kourelis, Adeline Harant, Hsuan Pain, and Joe Win (The Sainsbury Laboratory) for valuable discussions; Ioanna Morianou and Sophien Kamoun (The Sainsbury Laboratory) for reading the manuscript; Phil Robinson (John Innes Centre, Scientific Photographer), Damian Alger and Sophie Able (John Innes Centre, Horticultural Services) for their support.

## References

Baguma JK, Mukasa SB, Nuwamanya E, Alicai T, Omongo C, Hyde PT, Setter TL, Ochwo-Ssemakula M, Esuma W, Kanaabi M, et al (2023) Flowering and fruit-set in cassava under extended red-light photoperiod supplemented with plant-growth regulators and pruning. BMC Plant Biol 23: 335

Bally J, Jung H, Mortimer C, Naim F, Philips JG, Hellens R, Bombarely A, Goodin MM, Waterhouse PM (2018) The Rise and Rise of Nicotiana benthamiana: A Plant for All Reasons. Annu Rev Phytopathol 56: 405–426

Byrne M, Yeates DK, Joseph L, Kearney M, Bowler J, Williams MAJ, Cooper S, Donnellan SC, Keogh JS, Leys R, et al (2008) Birth of a biome: insights into the assembly and maintenance of the Australian arid zone biota. Mol Ecol 17: 4398–4417

Derevnina L, Kamoun S, Wu C-H (2019) Dude, where is my mutant? Nicotiana benthamiana meets forward genetics. New Phytol 221: 607–610

Ghosh S, Watson A, Gonzalez-Navarro OE, Ramirez-Gonzalez RH, Yanes L, Mendoza-Suárez M, Simmonds J, Wells R, Rayner T, Green P, et al (2018) Speed breeding in growth chambers and glasshouses for crop breeding and model plant research. Nat Protoc 13: 2944–2963

Hickey LT, N Hafeez A, Robinson H, Jackson SA, Leal-Bertioli SCM, Tester M, Gao C, Godwin ID, Hayes BJ, Wulff BBH (2019) Breeding crops to feed 10 billion. Nat Biotechnol 37: 744–754

Kourelis J, Kaschani F, Grosse-Holz FM, Homma F, Kaiser M, van der Hoorn RAL (2019) A homology-guided, genome-based proteome for improved proteomics in the alloploid Nicotiana benthamiana. BMC Genomics 20: 722

Lüdke D (2024) danlue/Speedy-benthi: Accelerating Nicotiana benthamiana generation times through extended photoperiods. doi: 10.5281/zenodo.10514569

Ranawaka B, An J, Lorenc MT, Jung H, Sulli M, Aprea G, Roden S, Llaca V, Hayashi S, Asadyar L, et al (2023) A multi-omic Nicotiana benthamiana resource for fundamental research and biotechnology. Nature Plants 9: 1558–1571

Schiavinato M, Marcet-Houben M, Dohm JC, Gabaldón T, Himmelbauer H (2020) Parental origin of the allotetraploid tobacco Nicotiana benthamiana. Plant J 102: 541–554

Schilling S, Melzer R, Dowling CA, Shi J, Muldoon S, McCabe PF (2023) A protocol for rapid generation cycling (speed breeding) of hemp (Cannabis sativa) for research and agriculture. Plant J 113: 437–445

Varshney RK, Bohra A, Roorkiwal M, Barmukh R, Cowling WA, Chitikineni A, Lam H-M, Hickey LT, Croser JS, Bayer PE, et al (2021) Fast-forward breeding for a food-secure world. Trends Genet 37: 1124–1136

Vollheyde K, Dudley QM, Yang T, Oz MT, Mancinotti D, Fedi MO, Heavens D, Linsmith G, Chhetry M, Smedley MA, et al (2023) An improved Nicotiana benthamiana bioproduction chassis provides novel insights into nicotine biosynthesis. New Phytol 240: 302–317

Watson A, Ghosh S, Williams MJ, Cuddy WS, Simmonds J, Rey M-D, Asyraf Md Hatta M, Hinchliffe A, Steed A, Reynolds D, et al (2018) Speed breeding is a powerful tool to accelerate crop research and breeding. Nature Plants 4: 23–29

