## Supplemental Data 2 for "Speedy benthi: Elongated photoperiods reduce generation times of the model plant species *Nicotiana benthamiana*"

Mean dpp for each condition:

condition	dpp

1	GH	82.75000

2	NB	78.75949

3	SB	67.65432

Standard deviation of the mean dpp for each condition:

condition	dpp

1	GH	1.251315

2	NB	2.760598

3	SB	2.134013

Shapiro-Wilk normality test for each condition:

data: data$dpp[data$condition == "NB"]

W = 0.9421, p-value = 0.001356

data: data$dpp[data$condition == "SB"]

W = 0.95671, p-value = 0.007974

data: data$dpp[data$condition == "GH"]

W = 0.88379, p-value = 0.02072

Kruskal-Wallis rank sum test comparing the conditions:

data: dpp by condition

Kruskal-Wallis chi-squared = 142.86, df = 2, p-value < 2.2e-16

Pairwise post-hoc test for each condition:

posthoc_dunn <- dunn.test(data$dpp, data$condition, method = "bonferroni")

Comparison of x by group

(Bonferroni)

Col Mean-|

Row Mean |	GH	NB

---------+-----------------------------------------------

NB |	3.052954

| 0.0034*

|

SB |	9.376256	9.972817

| 0.0000* 0.0000*

alpha = 0.05

Reject Ho if p <= alpha/2

Mean height for each condition:

condition	height

1	GH	98.62500

2	NB	58.36709

3	SB	61.70370

Standard deviation of the mean height for each condition:

condition	height

1	GH	6.045605

2	NB	2.639334

3	SB	2.852387

Shapiro-Wilk normality test for each condition:

data: data$height[data$condition == "NB"]

W = 0.95888, p-value = 0.01225

data: data$height[data$condition == "SB"]

W = 0.96908, p-value = 0.0472

data: data$height[data$condition == "GH"]

W = 0.95312, p-value = 0.417

Kruskal-Wallis rank sum test comparing the conditions:

data: height by condition

Kruskal-Wallis chi-squared = 86.12, df = 2, p-value < 2.2e-16

Pairwise post-hoc test for each condition:

> posthoc_dunn_height <- dunn.test(data$height, data$condition, method = "bonferroni")

Comparison of x by group

(Bonferroni)

Col Mean-|

Row Mean |	GH	NB

---------+------------------------------------------------

NB |	8.745920

| 0.0000*

|

SB |	5.134597	-5.737077

| 0.0000* 0.0000*

alpha = 0.05

Reject Ho if p <= alpha/2
