## Supplemental Data 5 for "Speedy benthi: Elongated photoperiods reduce generation times of the model plant species *Nicotiana benthamiana*"

Mean dpp for each condition:

condition	dpp

1	CN 72.35

2	CR 69.85

3	FN 80.55

4	FR 70.45

Standard deviation of the mean dpp for each condition:

condition	dpp

1	CN 2.207046

2	CR 2.896063

3	FN 2.605157

4	FR 2.297713

Shapiro-Wilk normality test for each condition:

data: data$dpp[data$condition == "FN"]

W = 0.82246, p-value = 0.001906

data: data$dpp[data$condition == "CN"]

W = 0.9594, p-value = 0.5319

data: data$dpp[data$condition == "FR"]

W = 0.96062, p-value = 0.1759

data: data$dpp[data$condition == "CR"]

W = 0.9633, p-value = 0.2168

Kruskal-Wallis rank sum test comparing the conditions:

data: dpp by condition

Kruskal-Wallis chi-squared = 57.923, df = 3, p-value = 1.632e-12

Pairwise post-hoc test for each condition:

> posthoc_dunn <- dunn.test(data$dpp, data$condition, method = "bonferroni")

Comparison of x by group

(Bonferroni)

Col Mean-|

Row Mean |	CN	CR	FN

---------+---------------------------------

CR |	2.837493

| 0.0136*

|

FN | -3.725941 -7.139839

| 0.0006* 0.0000*

|

FR |	2.174884 -0.811526	6.477231

| 0.0889 1.0000 0.0000*

alpha = 0.05

Reject Ho if p <= alpha/2

Mean height for each condition:

condition height

1	CN 59.1000

2	CR 62.1125

3	FN 64.3750

4	FR 67.5875

Standard deviation of the mean height for each condition:

condition	height

1	CN 1.909808

2	CR 2.202817

3	FN 2.723557

4	FR 1.849766

Shapiro-Wilk normality test for each condition:

data: data$height[data$condition == "FN"]

W = 0.94611, p-value = 0.3119

data: data$height[data$condition == "CN"]

W = 0.9764, p-value = 0.8798

data: data$height[data$condition == "FR"]

W = 0.95785, p-value = 0.1414

data: data$height[data$condition == "CR"]

W = 0.97157, p-value = 0.403

Levene's Test for Homogeneity of Variance (center = median):

Df F value Pr(>F)

group	3 1.2737 0.2867

116

Anova comparing the conditions:

aov(formula = height ~ condition, data = data)

Terms:

condition Residuals

Sum of Squares	1136.106	532.925

Deg. of Freedom	3	116

Residual standard error: 2.143404

Estimated effects may be unbalanced

> # Display the summary of ANOVA

> summary(anova_result)

Df Sum Sq Mean Sq F value Pr(>F)

condition	3 1136.1	378.7	82.43 <2e-16 ***

Residuals	116 532.9	4.6

---

Signif. codes: 0 ‘***’ 0.001 ‘**’ 0.01 ‘*’ 0.05 ‘.’ 0.1 ‘ ’ 1

Pairwise post-hoc Tukey multiple comparisons of means:

95% family-wise confidence level

Fit: aov(formula = height ~ condition, data = data)

$condition

diff	lwr	upr	p adj

CR-CN 3.0125 1.4823997 4.542600 0.0000069

FN-CN 5.2750 3.5081924 7.041808 0.0000000

FR-CN 8.4875 6.9573997 10.017600 0.0000000

FN-CR 2.2625 0.7323997 3.792600 0.0010822

FR-CR 5.4750 4.2256783 6.724322 0.0000000

FR-FN 3.2125 1.6823997 4.742600 0.0000015
